# Midkine Dependency Defines a Therapeutically Actionable Vulnerability in Medulloblastoma

**DOI:** 10.64898/2026.04.11.717918

**Authors:** Sridharan Jayamohan, Prabhakar P. Venkata, Jessica D. Johnson, Baskaran Subramani, Nour Abdelfattah, Yi Zou, Zhao Lai, Shayla A. Hernandez, Panneerdoss Subbarayalu, Uday Pratap, Siyuan Zheng, Manjeet K. Rao, Andrew J. Brenner, Suryavathi Viswanadhapalli, Ratna K. Vadlamudi, Kyuson Yun, Hareesh B. Nair, Gangadhara R. Sareddy

## Abstract

Medulloblastoma (MB), the most common malignant pediatric brain tumor, remains associated with high recurrence rates and substantial treatment-related morbidity despite aggressive multimodal therapy. Here, we identify midkine (MDK), a secreted growth factor/cytokine, as a previously unrecognized central regulator of MB growth and survival. Tissue microarray analysis revealed elevated MDK protein expression in MB specimens, and single-cell RNA sequencing datasets showed tumor cell-specific MDK expression. Genetic depletion or antibody-mediated neutralization of MDK suppressed proliferation and survival and induced apoptosis across multiple MB cell lines, whereas recombinant MDK supplementation exerted the opposite effect. Mechanistically, phosphokinase profiling and immunoblot analyses showed that MDK signals through multiple receptors, including nucleolin (NCL), LRP1, and syndecan-2 (SDC2), to sustain oncogenic ERK1/2 and Akt–mTOR–S6 signaling. Transcriptomic profiling following MDK silencing or depletion revealed marked suppression of ribosome biogenesis, global protein translation, and MYC-driven programs, coupled with activation of inflammatory and apoptotic responses. Consistently, mechanistic studies utilizing RPS6 staining, ribosomal RNA quantification, puromycin SUnSET, and OPP incorporation assays confirmed that loss of MDK impairs ribosome biogenesis and protein synthesis. Notably, MDK suppression also triggered robust activation of the IFN–cGAS–STING pathway, linking translational stress to innate immune signaling. In orthotopic xenograft models, CRISPR-mediated MDK knockout significantly reduced tumor growth and prolonged survival. Together, these findings establish MDK as a key integrator of oncogenic signaling, translational control, and innate immunity in MB, highlighting MDK as a compelling therapeutic target.

## Introduction

Medulloblastoma (MB) is the most common malignant pediatric brain tumor, accounting for ∼20% of childhood brain tumors and occurring at an annual incidence of approximately 5 cases per million children[1, 2]. MB is a biologically heterogeneous disease composed of distinct molecular subgroups. The 2021 WHO Classification of Central Nervous System Tumors delineates four genetically defined subgroups: WNT-activated, SHH-activated/TP53-wildtype, SHH-activated/TP53-mutant, and non-WNT/non-SHH (formerly referred to as Groups 3 and 4)[3], with further refinement into four SHH subtypes and eight non-WNT/non-SHH subtypes[4–6]. The current standard of care for MB involves maximal safe surgical resection followed by adjuvant multi-agent chemotherapy, except in infants, risk-adapted craniospinal irradiation[1, 2]. Although these multimodal therapies have improved overall 5-year survival, outcomes remain poor for high-risk patients, including those under 3 years of age, individuals with residual or metastatic disease, and patients classified within molecularly aggressive subgroups[7]. Moreover, long-term survivors often experience substantial treatment-related toxicities, including neurocognitive impairment, endocrine dysfunction, and reduced quality of life[8]. Therapy resistance and recurrence further remain major clinical challenges[7] underscoring the urgent need for more effective and less toxic therapeutic strategies for MB. Despite advances in molecular classification, the key signaling mechanisms driving MB pathogenesis remain incompletely understood, highlighting the critical need to identify novel molecular vulnerabilities and therapeutic targets.

Midkine (MDK), is a multifunctional secreted protein that functions as both a cytokine and a growth factor, modulating numerous intracellular signaling pathways[9–11]. The biological effects of MDK are highly tissue- and cell-specific and are determined by the repertoire of MDK receptors expressed on target cells. Several receptors have been identified, including anaplastic lymphoma kinase (ALK)[12], protein tyrosine phosphatase ζ (PTPζ/PTPRZ1)[13], Notch2[14], low-density lipoprotein (LDL)-receptor-related protein (LRP1)[15], integrins (α6β1 and α4β1)[16], syndecan (SDC2) [12], and nucleolin (NCL)[17, 18]. Through these interactions, MDK activates oncogenic kinase signaling and regulates transcription factors including STAT3, Hes-1, β-catenin, and NF-κB. MDK is overexpressed across multiple cancer types and is recognized as both a diagnostic and prognostic biomarker associated with poor survival[19, 20]. Functionally, MDK contributes to numerous hallmarks of cancer, including enhanced cell proliferation, survival, evasion of apoptosis, migration, invasion, metastasis, angiogenesis, and modulation of antitumor immunity and inflammation[20]. Physiologically, MDK is highly expressed during embryonic and fetal development, with markedly reduced expression in postnatal tissues[21]. Notably, serial analysis of gene expression (SAGE) studies identified MDK as one of the most highly expressed transcripts in MB tissues, with little to no expression in normal cerebellum or other pediatric and adult brain regions[22]. However, the mechanistic role of MDK in driving MB progression remains poorly defined, representing a significant gap in our understanding of MB biology.

In this study, we demonstrated the functional significance of MDK in MB pathogenesis. We demonstrate that MDK is highly expressed in MB tissues. Genetic silencing or MDK-antibody-mediated neutralization showed that MDK is essential for MB cell growth and survival. Comprehensive transcriptomic profiling and mechanistic studies revealed that MDK signals through the MAPK and AKT/mTOR/S6 axis to promote MB growth by enhancing ribosome biogenesis and global protein translation while suppressing cytotoxic interferon signaling pathways. Collectively, our findings identify MDK as a critical driver of MB biology and provide compelling evidence that targeting MDK may represent a viable and innovative therapeutic strategy for patients with MB.

## Materials and Methods

### Cell lines and Reagents

Daoy, D556, D283-Med medulloblastoma (MB) cell lines were obtained from the American Type Culture Collection (ATCC, Manassas, VA) and maintained according to ATCC-recommended culture conditions. HD-MB03 cells were kindly provided by Dr. Till Milde[23], German Cancer Research Center (DKFZ). All cell lines were passaged fewer than 15 times after receipt or resuscitation. Mycoplasma contamination was routinely assessed using the Mycoplasma PCR Detection Kit (Sigma-Aldrich, St. Louis, MO), and all cell lines tested negative. Cell identity was confirmed by short tandem repeat (STR) profiling. All procedures complied with the Helsinki Declaration and institutional guidelines approved by the UT Health San Antonio Institutional Review Board. CellTiter-Glo Luminescent Cell Viability Assay and luciferase assay reagents were obtained from Promega (Madison, WI). Annexin V/PI kits were purchased from BioLegend (San Diego, CA). Antibodies against p-Akt (S473), Akt, p-mTOR (S2448), mTOR, p-S6 (S235/236), S6, p-ERK (Thr202/Tyr204), ERK, pRSK1/2/3, p-4E-BP1 (Thr37/46), 4E-BP1, STING, p-STING (S366), TBK1/NAK, p-TBK/NAK (S172), cGAS, β-actin and GAPDH were obtained from Cell Signaling Technology (Beverly, MA). Ki67 antibody was purchased from Abcam (Waltham, MA). Midkine (MDK) polyclonal antibodies were purchased from Proteintech (11009-1-AP) and Santa Cruz Biotechnology (Dallas, TX). C-Myc antibody was purchased from Santa Cruz Biotechnology (Dallas, TX).

### MDK siRNA Transfection and CRISPR gRNA Transduction

MB cells were transfected with scrambled or MDK-specific siRNAs using Lipofectamine RNAiMAX (Thermo Fisher; Cat# 13778150) according to the manufacturer’s protocol. Cells were collected 48–72 h post-transfection for downstream analyses (Western blotting, viability, colony formation, and Annexin V assays). The following siRNAs (Millipore Sigma) were used: Universal Negative Control #1; MDK siRNA 1 (SASI_Hs01_00094499); MDK siRNA 2 (SASI_Hs01_00094501); MDK siRNA 3 (SASI_Hs01_00094503). Daoy MDK-knockout (KO) cells were generated using human MDK CRISPR gRNA lentiviral particles (HSPD0000025378). Cells were selected using puromycin (1 µg/mL). Lentiviral particles expressing non-targeting gRNA were used as controls.

### Cell viability, Clonogenic, and Apoptosis assays

Cell viability was assessed using the CellTiter-Glo 2.0 Luminescent Assay (Promega; Cat# G9241) according to the manufacturer’s instructions, and luminescence was measured using a Promega GloMax® luminometer. Clonogenic assays were performed in 6- or 12-well plates as previously described. Apoptosis was assessed using an Annexin V/PI staining kit (BioLegend) following previously published methods.

### RNA-sequencing and RT-qPCR

RNA-seq was performed as previously described[24]. Daoy cells were transfected with two independent MDK siRNAs (siRNA1 and siRNA2); D283 Med cells were transfected with MDK siRNA1. RNA was isolated 72 h post-transfection using the RNeasy Mini Kit (Qiagen). For antibody studies, Daoy cells were treated with IgG or MDK-neutralizing antibody (5 µg/mL) for 48 h prior to RNA isolation. Libraries were prepared using the Illumina TruSeq Stranded mRNA Kit and sequenced on an Illumina HiSeq 3000 platform at the UTHSA Genome Sequencing Facility. Differential gene expression analysis was performed using DESeq2; pathway analysis was conducted using Gene Set Enrichment Analysis (GSEA). GEO submission is pending. For RT-qPCR, cDNA was synthesized using the Applied Biosystems cDNA Synthesis Kit, and quantitative PCR was performed using SYBR Green master mix and gene-specific primers from the Harvard Primer Bank (http://pga.mgh.harvard.edu/primerbank/).

### Cell lysis and Western Blotting

Cells were lysed in RIPA buffer (Thermo Scientific; Cat# PI89901) supplemented with EDTA and Halt Protease/Phosphatase Inhibitor Cocktail (Thermo Scientific; Cat# PI78441) for 30 min at 4°C with gentle rotation. Lysates were resolved by SDS-PAGE and transferred onto Nitrocellulose membranes (Bio-Rad; Cat# 1620115). Membranes were blocked in 5% non-fat milk in TBST for 1 h, incubated with primary antibodies overnight at 4°C, washed, and probed with secondary antibodies for 1 h at room temperature: Anti-mouse IgG HRP (CST; Cat# 70765; 1:1000); Anti-rabbit IgG HRP (Cytiva; Cat# NA9341; 1:2000). Signal was detected using enhanced chemiluminescent substrate (Millipore Sigma; Cat# WBKLS0500) and imaged using a Bio-Rad ChemiDoc MP system.

### MDK ELISA assay

The Human MDK ELISA kit (Invitrogen; Cat#EH319RB) was used to quantify the level of secreted MDK in MB cells, and HA cells used as a control. Briefly, 1X10^6^ cells were seeded in a culture medium. After 12h, the culture medium was removed, and the cells were cultured in serum-free medium for 48h. Following this incubation period, the conditioned medium was collected, and the assay was performed according to the manufacturer’s instructions.

### Immunofluorescence

MB cells treated with control or MDK-specific siRNA, or with IgG or MDK-neutralizing antibody (2.5µg/ml), were fixed with ice-cold methanol for 20 min and permeabilized with 0.2% Triton X-100. After blocking with normal goat serum, cells were incubated with primary cGAS antibody (1:200) overnight at 4°C, followed by Alexa Fluor-conjugated secondary antibodies (Alexa 594; Thermo Fisher). Coverslips were mounted with ProLong Gold Antifade with DAPI (Invitrogen; Cat# P36980). Images were captured using a Zeiss confocal microscope.

### Tissue Microarrays and Immunohistochemistry

MB tissue microarrays (TMAs) were obtained from TissueArray.com (Derwood, MD), containing 20 MB cases (60 cores) and 3 normal brain tissues. TMA processing and MDK staining were performed as previously described. In brief, slides were deparaffinized, rehydrated, subjected to citrate-based antigen retrieval (Vector Labs), treated with 3% H₂O₂, blocked, and incubated overnight with MDK antibody. Detection was visualized using DAB substrate and hematoxylin counterstain. For mouse experiments, orthotopic xenograft sections were stained for Ki67 and pS6.

### Ribosomal Biogenesis

Ribosome biogenesis was evaluated by RPS6 immunofluorescence. Cells were treated with MDK siRNA or MDK antibody for 24 h, fixed, permeabilized, blocked, and incubated with anti-RPS6 antibody (1:100; Santa Cruz; Cat# sc-74459), followed by Alexa 488 secondary antibody. Five fields per condition were imaged, and mean fluorescence intensity for RPS6 was quantified using ImageJ (Java 1.8.0; NIH).

### O-Propargyl-puromycin (OPP) and Puromycin SUnSET assays

Cayman’s Protein Synthesis Assay Kit (Cat# 601100; Cayman Chemical, Ann Arbor, MI, USA) was used to measure protein synthesis according to the manufacturer’s instructions. Briefly, MB cells were plated in 8-well chamber slides, treated with MDK-neutralizing antibody for 24 h, and then incubated with the cell-permeable, alkyne-containing puromycin analog O-propargyl-puromycin (OPP; 2.5 μL/mL) for 90 min at 37°C. Cells were processed for fluorescence detection following the kit protocol, and 5-FAM–based OPP incorporation was visualized by confocal microscopy. Five fields per condition were imaged, and mean fluorescence intensity for OPP was quantified using ImageJ (Java 1.8.0; NIH). Values were calculated by dividing total fluorescence intensity by the number of cells in each field and normalized to the respective control group. Global protein synthesis was further assessed using the puromycin-based SUnSET (Surface Sensing of Translation) method. MB cells were transfected with MDK siRNA or treated with MDK antibody (10, 20 μg/mL) for 16 h and subsequently incubated with puromycin (1 μM; P7255, Millipore Sigma) for 30 min. Total cell lysates were collected and immunoblotted using the anti-puromycin antibody [3RH11] (Kerafast; Cat# EQ0001). Cycloheximide (CHX)-treated cells served as positive controls for inhibition of protein synthesis.

### Bulk Transcriptomic Analysis of Human Medulloblastoma

To evaluate MDK expression across pediatric brain tumors, we used the Gliovis (gliovis.bioinfo.cnio.es) web tool, which included the Griesinger et al., data set[25] (n=130; non-tumor=13, medulloblastoma=22, ependymoma=46, pilocytic astrocytoma=15 and glioblastoma=34). To evaluate MDK expression across human medulloblastoma (MB) subgroups, we analyzed the publicly available Cavalli et al [26] bulk microarray dataset (GEO: GSE85217). Log2-transformed MDK mRNA expression values were compared across the four consensus molecular subgroups (WNT, SHH, Group 3, and Group 4). Statistical significance between groups was determined using analysis of variance (ANOVA). Data were visualized as box plots using the ggplot2 R package (v3.4.4; RRID:SCR_014601).

### Single-Cell RNA Sequencing (scRNA-seq) Processing and Analysis

External scRNA-seq data from the Riemondy et al[27]. Human MB cohort (GEO: GSE155446) were obtained as count matrices and processed utilizing the Seurat R package (v5.2.1; RRID:SCR_016341). Single-cell gene expression counts were normalized to library size and log2-transformed. Principal component analysis (PCA) was performed using the top 2,000 most variable genes, followed by batch correction using Harmony (v1.2.0; RRID:SCR_018809). Dimensionality reduction was applied to generate spatial feature plots illustrating MDK expression across subgroups. Furthermore, cellular annotations were used to generate violin plots comparing MDK expression levels specifically between malignant cells and the immune/stromal compartments.

### Intercellular Communication Analysis

To infer intercellular signaling networks within the MB tumor microenvironment, we utilized the CellChat R package (v2.2.1; RRID:SCR_021946) on the processed scRNA-seq dataset. CellChat’s internal, literature-validated database of ligand-receptor interactions was used to construct communication networks. Analysis was specifically targeted to the Midkine (MK) signaling pathway to characterize the directionality and interaction strength between malignant tumor cells (the primary ligand source) and stromal components.

### In vivo Orthotopic Tumor Model

All animal work was approved by the UT Health San Antonio IACUC. NOD.CB17-Prkdcscid/NCrCrl mice (8–10 weeks, Charles River) were used. Daoy cells expressing control gRNA or MDK-targeting gRNA were labeled with GFP-luciferase, and 1 × 10⁶ cells were injected orthotopically into the cerebellum (1.5 mm lateral, 1.5 mm posterior to lambda, 2.0 mm depth). Tumor growth was monitored weekly by bioluminescence imaging (Xenogen IVIS). Mice were euthanized when moribund or neurologically impaired. Survival was analyzed using Kaplan–Meier curves and log-rank testing.

### Statistical analysis

Statistical analyses were performed using GraphPad Prism 10 software. Comparisons between groups were made using Student’s t-test or one-way ANOVA where appropriate. Data are presented as mean ± SEM. Statistical significance was defined as p < 0.05.

## Results

### MDK is essential for MB cell growth and survival

MDK has been implicated in multiple oncogenic processes; however, its functional significance in MB has remained undefined. Analyses of pediatric brain tumor datasets (Gresinger et al[25]) revealed that MB exhibits the highest MDK expression relative to normal brain and other pediatric brain tumor types (**Fig. 1A**). Consistently, immunohistochemical analysis of MB tissue microarrays showed robust MDK protein expression across tumor samples, whereas normal cerebellum showed minimal staining (**Fig. 1B,C**). To further investigate the potential role of MDK in MB, we evaluated its transcriptomic expression across the four consensus molecular subgroups (WNT, SHH, Group 3, and Group 4). Analysis of the Cavalli et al[5] bulk microarray dataset revealed that MDK mRNA is abundantly expressed across all MB subgroups. Notably, we observed a significant variance in expression levels among the groups, with the WNT subgroup exhibiting the highest overall MDK expression followed by group 4 (**Fig. 1D**). To determine the cell-type-specific origins of MDK within the heterogeneous tumor microenvironment, we analyzed single-cell RNA sequencing (scRNA-seq) data from the Riemondy et al.,[28] MB cohort. Comparing expression between cellular compartments revealed that MDK is highly expressed in malignant cells across all four molecular subgroups, with minimal to no MDK expression detected in the immune stroma (**Fig. 1E,F**). Importantly, we confirmed MDK secretion by tumor cells using ELISA, which demonstrated that MB tumor cells secrete substantial levels of MDK, whereas astrocytes exhibited minimal MDK secretion (**Fig. 1G**). Collectively, these findings indicate that MB tumor cells are the primary source of MDK and that MDK functions as an autocrine growth factor to sustain tumor cell proliferation.

**Fig. 1.**
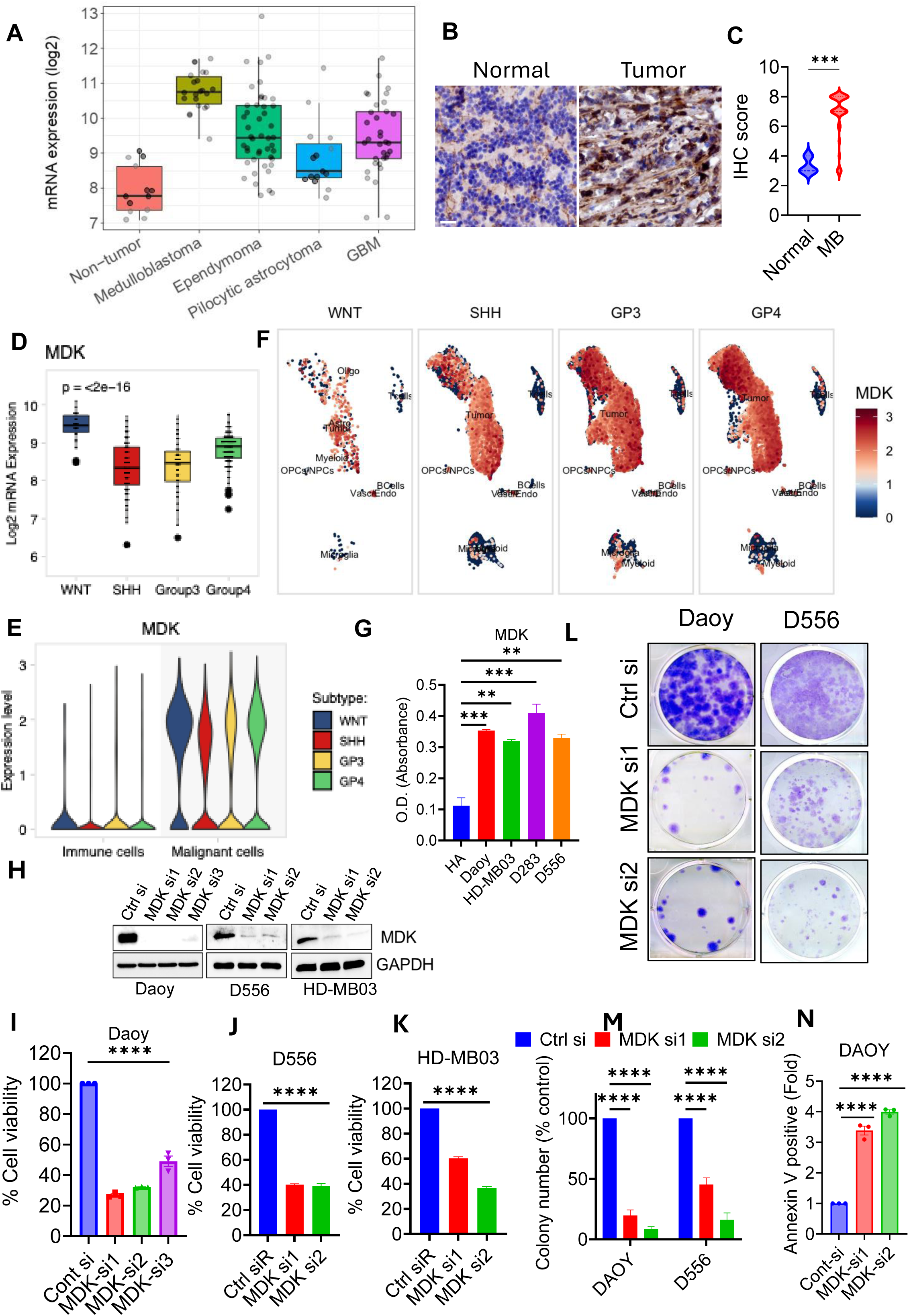
MDK is highly expressed in MB and is essential for MB cell proliferation and survival. A, Boxplots of MDK gene expression in pediatric brain tumor datasets (Griesinger et al., data sets[25]). B–C, MB tissue microarray (normal = 3, MB = 60) subjected to IHC analysis to determine MDK expression (B) and quantification (C). D, Box plot illustrating MDK log2 mRNA expression across four medulloblastoma (MB) subgroups (WNT, SHH, Group 3, and Group 4). Data was analyzed from the Cavalli et al. human MB microarray dataset (GEO: GSE85217). E, Violin plots comparing MDK expression levels between immune cells (stromal) and malignant cells across the four MB subtypes (WNT, SHH, GP3, GP4). Data is derived from a meta-analysis of the Reimondy et al[28] single-cell RNA sequencing (scRNA-seq) dataset (GEO: GSE155446). F, Single-cell feature plots divided by MB subgroup showing the spatial distribution of MDK expression. Color gradients indicate expression levels (red = high, blue = low). G, Secreted levels of MDK protein were determined in conditioned media collected from human astrocytes (HA) and different MB cells using the MDK ELISA assay. H, MDK expression was silenced in Daoy, D556, and HD-MB03 MB cells using different MDK-siRNAs, and knockdown was confirmed by Western blotting. I-K, Effect of MDK silencing on the cell viability of Daoy, D556, and HD-MB03 cells determined using CellTiter-Glo assays (5 days). L-M, Effect of MDK silencing on the clonogenic survival was assessed using colony-formation assays (2 weeks), and colony number was quantified. N, Effect of MDK silencing on the apoptosis was measured using Annexin V assays (72 h). **p<0.01, ***p < 0.001; ****p < 0.0001. ANOVA

To evaluate MDK functional relevance, we silenced MDK using multiple independent siRNAs (**Fig. 1H**) and assessed its impact on MB cell proliferation, survival and apoptosis. MDK knockdown significantly reduced proliferation (**Fig. 1I-K**), impaired clonogenic survival (**Fig. 1L,M**), and induced apoptosis (**Fig. 1N**) across multiple MB cell lines. Since MDK is a secreted protein, we neutralized extracellular MDK using an MDK-blocking antibody and assessed its effect on MB cells. Therapeutic targeting of MDK with MDK-neutralizing antibody phenocopied these effects, suppressing proliferation and colony formation while promoting apoptotic cell death (**Fig. 2A-D**) across multiple MB cell lines. Together, these findings establish MDK as an essential survival factor in MB, supporting its role as a therapeutically actionable oncogenic driver.

**Fig. 2.**
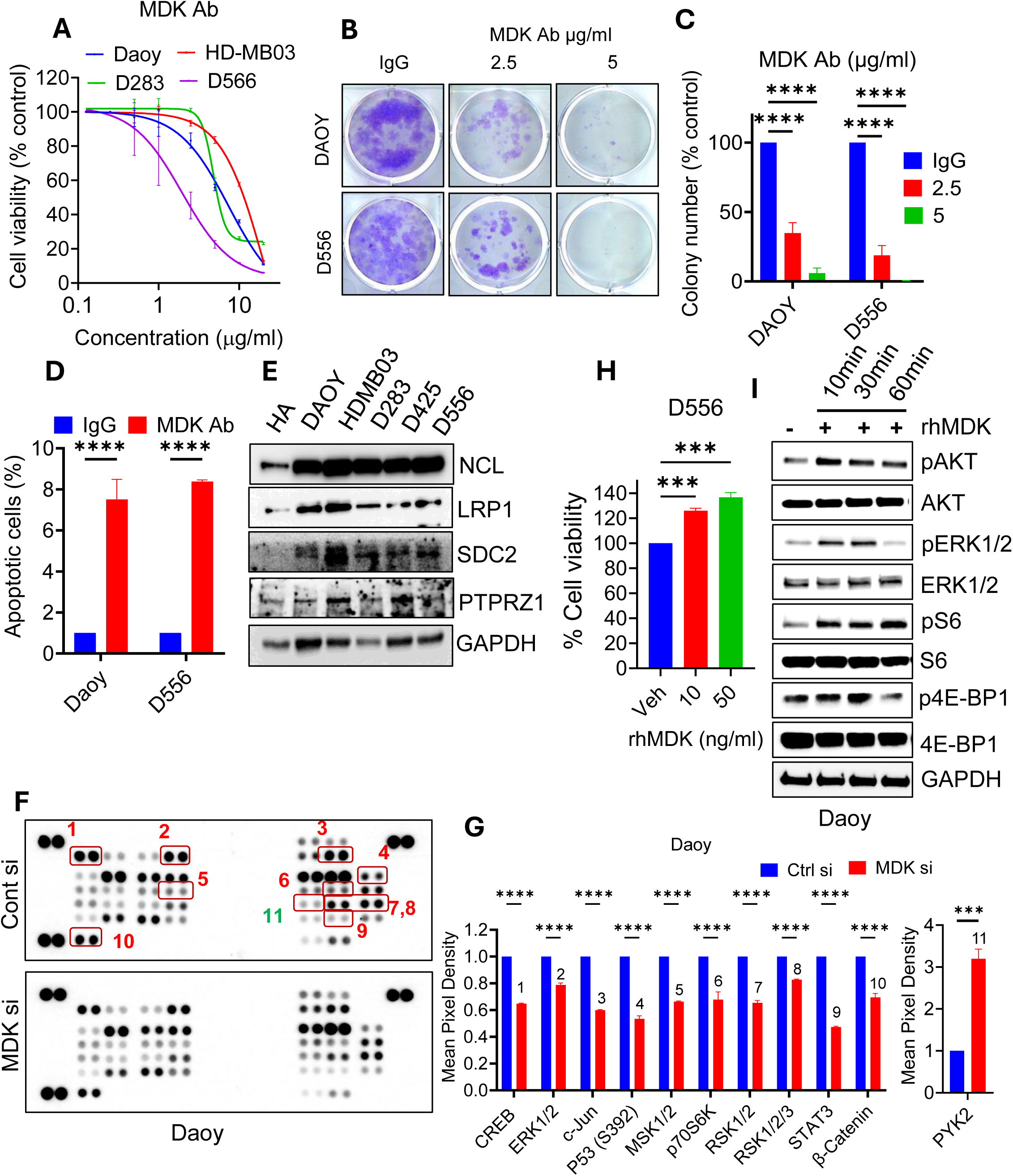
MDK antibody treatment suppressed MB cell growth, and MDK promotes oncogenic kinase signaling. A, MB cells were treated with either control IgG or MDK-antagonizing antibody (MDK-Ab; 4 days), and effects on viability were determined using CellTiter-Glo assays. B-C, Cell survival following MDK-Ab treatment was evaluated using clonogenic assays (2 weeks) and colony number was quantified. D, Apoptosis was determined using Annexin V assays following MDK-Ab treatment (48 h). E, Expression levels of known MDK receptors in human astrocytes (HA) and MB cells were determined using western blotting. F-G, MDK was silenced using siRNA and lysates from control and MDK-KD cells were subjected to R&D Phospho-Kinase Arrays. Quantification of array band intensities was shown. H, D556 MB cells were cultured with and without supplementation of recombinant MDK protein (rMDK) for 48h, and cell proliferation was determined using Cell-Titer-Glo assays. I, Daoy cells were stimulated with rMDK protein for the indicated time points, and its effect on kinase signaling was determined using Western blotting for the indicated proteins. ***p < 0.001; ****p < 0.0001. ANOVA

### MDK suppression downregulates oncogenic kinase signaling pathways

MDK is known to signal through several cell-surface receptors, including ALK, PTPRZ1, NCL, LRP1, integrins, and SDC2, and activate oncogenic kinase signaling cascades to drive abnormal cell growth[29]. Recent cell–cell communication analyses in MB tissues suggest that NCL and PTPRZ1 are dominant MDK receptors[30] ; however, their functional roles in MB remain unclear. Western blot analysis confirmed that MB cells express PTPRZ1, NCL, LRP1, Notch2, and SDC2, with notably high levels of NCL (**Fig. 2E**). To delineate downstream kinase signaling, we performed phosphokinase array profiling on MDK-silenced cells. MDK knockdown decreased the phosphorylation of several oncogenic kinases, including ERK1/2, RSK1/2, p70S6K, and STAT3, key regulators of proliferation, survival, and resistance to apoptosis (**Fig. 2F,G**). We further examined whether MDK protein functions as a growth factor in MB cells by supplementing with recombinant MDK protein (rMDK). rMDK supplementation significantly increased the proliferation of MB cells (**Fig. 2H**) and promoted the growth factor signaling, such as phosphorylation of MAPK and AKT signaling cascades (**Fig. 2I**). Conversely, Western blot analysis using two independent siRNAs and MDK-neutralizing antibody confirmed decreased activation of pERK1/2/pRSK1/2 signaling and Akt/mTOR/S6/4EBP1 signaling cascades in Daoy and D556 MB cells (**Fig. 3A-D**). Together, these findings indicate that MDK promotes MB growth by engaging multiple receptors to activate oncogenic kinase signaling pathways critical for cell proliferation and survival.

**Fig. 3.**
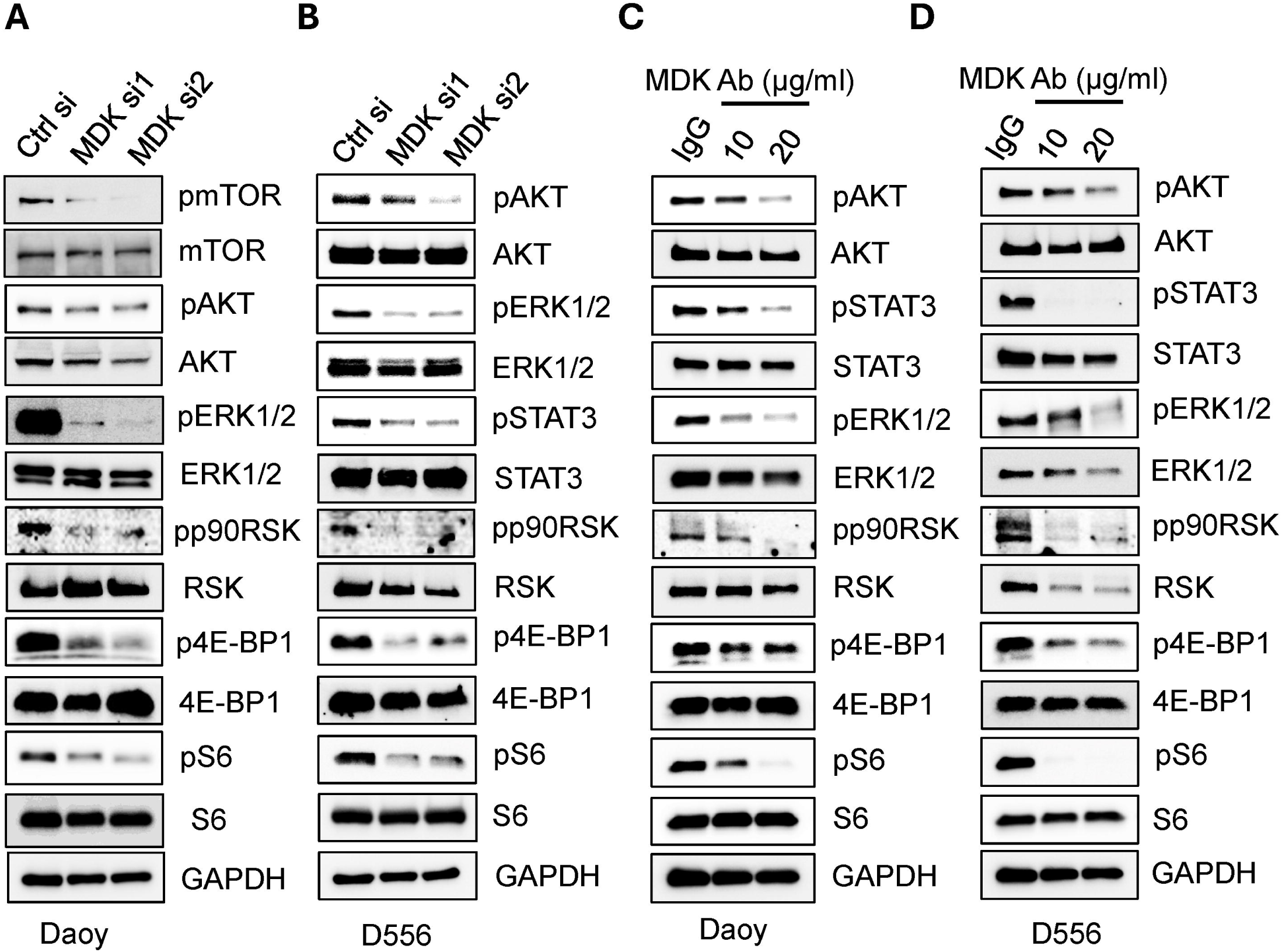
MDK silencing or neutralization attenuated oncogenic kinase signaling in MB cells. A-B, Daoy and D556 cells were transfected with two different siRNAs; after 48 h, lysates were subjected to Western blotting for the indicated proteins. C-D, Daoy and D556 cells were treated with control IgG or MDK antibody (10 or 20 µg; 24 h) and subjected to Western blotting for the indicated proteins.

### MDK silencing or antagonization alters gene expression programs governing translation and immune pathways

To elucidate the genomic changes driven by MDK signaling, we performed RNA-seq on Daoy cells transfected with two independent siRNAs. Gene set enrichment analysis (GSEA) showed that MDK silencing led to significant downregulation of ribosomal and translation pathways, MYC target genes, and mTOR signaling pathways, and upregulation of inflammatory response, interferon signaling, and apoptotic pathways across both MDK-siRNA datasets (**Fig. 4A-B**). Importantly, we confirmed whether MDK neutralization will have similar effects on gene expression. Treatment of MBs with an MDK-neutralizing antibody, recapitulated these results (**Fig. 4C**). Additionally, we validated these results in another MB cell line, D283, which also showed that MDK silencing led to negative enrichment of ribosomal and translation pathways, MYC targets, and mTOR signaling, along with positive enrichment of inflammatory and cell-death pathways (**Fig.4D**). GSEA plots and corresponding gene-expression heatmaps demonstrated consistent regulation of these pathways across different MB cell lines and under both siRNA and antibody treatment conditions (**Fig. 4E-G; Supplementary Fig. 1A**). Further validation in additional MB cells such as D556 and HD-MB03 using RT-qPCR confirmed that MDK suppression led to decreased expression of genes involved in ribosome biogenesis (**Fig. 4H-K**). Collectively, these findings establish that silencing MDK expression or treating cells with an MDK antibody significantly decreases the expression of genes involved in protein translation and ribosomal biogenesis, while its inhibition reprograms the transcriptome toward immune response signaling and apoptosis.

**Fig. 4.**
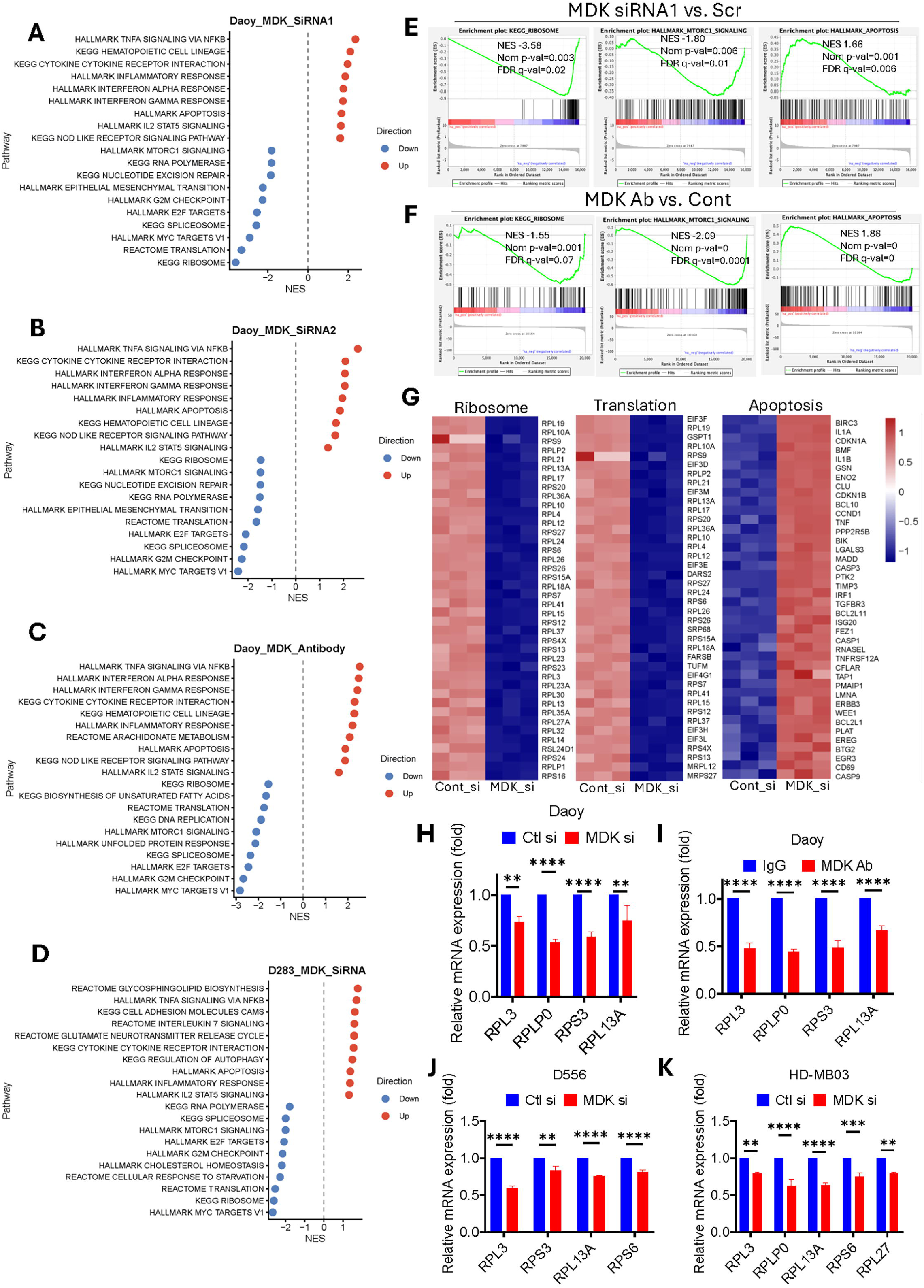
MDK regulates ribosome/translation and immune-response genes in MB cells. A–B, Daoy cells were transfected with control siRNA or MDK siRNA1 (A) or siRNA2 (B). RNA was isolated and subjected to RNA-seq and GSEA analysis; the top significantly enriched positive and negative pathways shared between the two siRNA groups are shown. C, Daoy cells treated with control IgG or MDK-antagonizing antibody (48 h) were subjected to RNA-seq and GSEA analysis. Top significantly enriched positive and negative pathways are shown. D, D283 cells were transfected with control or MDK siRNA1; RNA was isolated and subjected to RNA-seq and GSEA analysis. Top significantly enriched positive and negative pathways are shown. E–F, GSEA plots showing negative enrichment of ribosome and mTORC1 signaling and positive enrichment of apoptosis in MDK-siRNA (E) and MDK-Ab-treated (F) cells. G, Heatmap showing downregulation of ribosome/translation genes and upregulation of apoptosis-related genes. H–K, RT-qPCR demonstrating downregulation of ribosomal genes in Daoy MDK-siRNA (H), Daoy MDK-Ab-treated (I), D556 MDK-siRNA (J), and HD-MB03 MDK-siRNA (K) cells. **p < 0.01; ***p < 0.001; ****p < 0.0001. ANOVA

### MDK suppression decreases ribosomal biogenesis and global protein synthesis

Our transcriptomic analyses consistently showed that MDK suppression decreases the expression of mTORC1 signaling, MYC targets, and ribosome/translation pathway genes, and Western blot analyses demonstrated consistent downregulation of pERK1/2 and Akt/mTOR pathway components, including ribosomal proteins p90RSK1/2, pS6, and the translation factor p4EBP1, which are essential drivers of ribosomal biogenesis and protein synthesis. Dysregulated ribosome production and translation are hallmarks of cancer, and MB cells, like many tumor types, rely on elevated ribosome biogenesis to support rapid proliferation*[31],[32–34]*. Prior studies have shown that MYC and mTOR are key regulators of ribosome biogenesis and protein translation in MB[35–37]: MYC amplifies transcription of ribosomal components and rRNA, while mTOR activates the translational machinery through phosphorylation of 4E-BP1 and S6. Moreover, ERK1/2 and mTOR signaling enhance MYC stability and transcriptional activity through post-translational modifications and downstream signaling cascades[38–40]. Based on these observations, we hypothesized that MDK inhibition suppresses ribosome biogenesis and protein translation by disrupting MYC-driven transcriptional programs and the MAPK and mTOR signaling cascades. We first examined the role of MDK in regulating MYC protein expression using western blotting. As shown in **Fig. 5A**, MDK silencing or neutralization with an antibody resulted in decreased MYC expression in MB cells. To determine whether MDK blockade directly affects ribosomal biogenesis, we examined expression of the 40S ribosomal subunit protein rpS6 as a marker of ribosomal content. MB cells were transfected with MDK siRNA or treated with MDK antibody and stained for rpS6. Compared with controls, both MDK-silenced and MDK-antibody-treated cells showed reduced rpS6 immunofluorescence (**Fig. 5C-F**). Given that pre-rRNA processing and synthesis are critical rate-limiting steps in ribosome biogenesis, and since MYC and mTOR regulate rRNA transcription, we next evaluated rRNA synthesis following MDK inhibition. RT-qPCR analysis revealed that MDK knockdown or antibody treatment significantly decreased the levels of mature 5.8S and 28S rRNAs, as well as pre-rRNA (**Fig. 5G-H**).

**Fig. 5.**
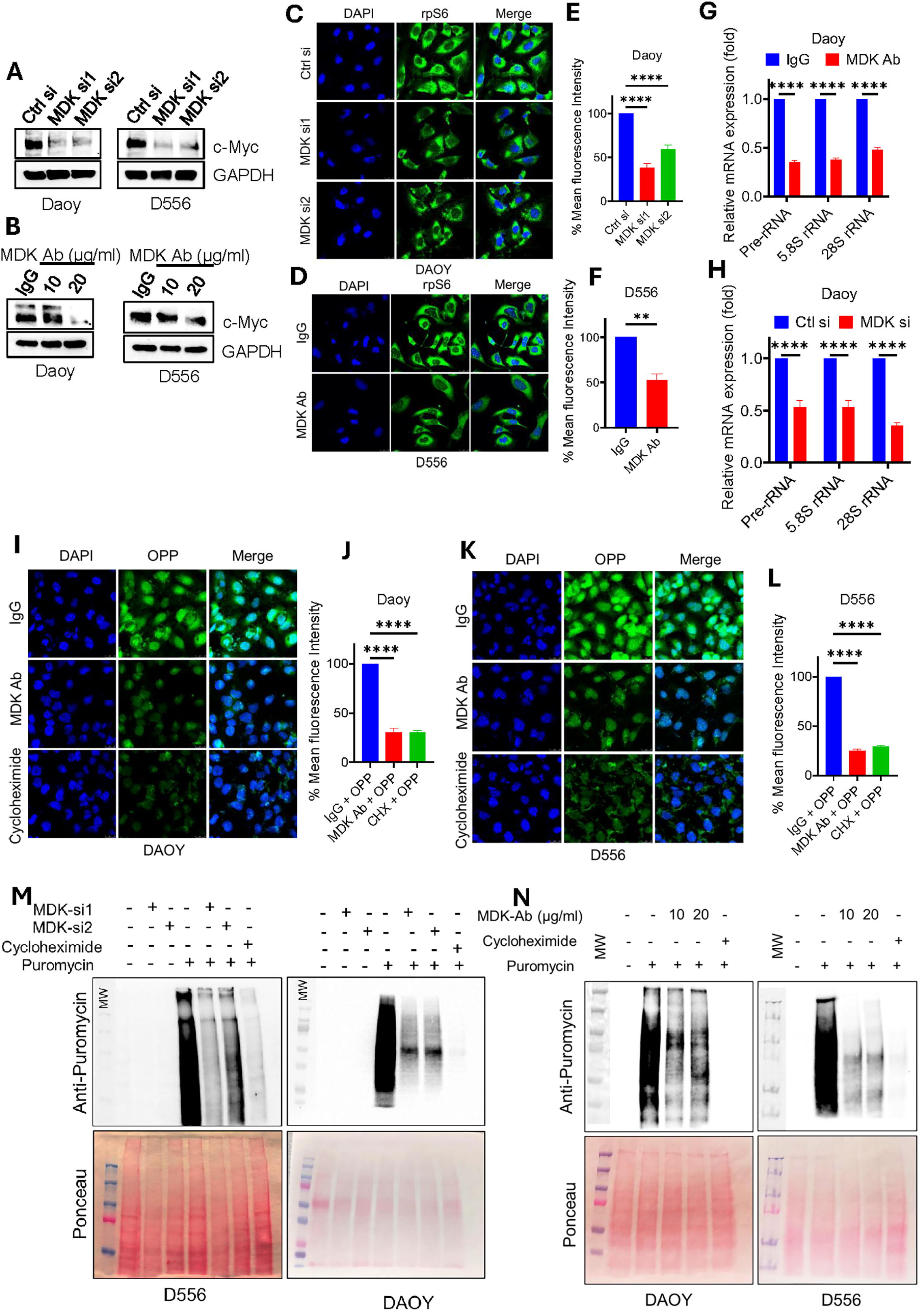
MDK suppression reduces ribosomal biogenesis and new protein synthesis. A-B, Daoy and D556 cells were treated with MDK-siRNA (A) or MDK-antibody (B), and c-Myc expression was determined using Western blotting. C-F, Daoy (C) and D556 (D) cells were fixed 72 h after MDK-siRNA transfection or MDK-Ab treatment (20 µg), and rpS6 (40S ribosomal subunit) was visualized using confocal microscopy. Quantification of total fluorescence intensity (E,F) was performed using ImageJ; mean fluorescence intensity normalized to control is shown. G-H, Daoy cells transfected with MDK-siRNA (E) or treated with MDK-Ab (5 µg; 48 h) (F) were analyzed for pre-rRNA, 5.8S, and 28S rRNA levels by RT-qPCR. I-L, Daoy and D556 cells were treated with MDK-Ab (20 µg; 24 h), followed by incubation with OPP (90 min), a puromycin analog incorporated into newly synthesized proteins; reduced FITC signal was visualized by confocal microscopy. Quantification of total fluorescence intensity was performed using ImageJ; mean fluorescence intensity normalized to control is shown. M-N, D556 and Daoy cells transfected with two independent MDK-siRNAs (M) or treated with MDK-Ab (10, 20 µg; 24 h) (N) were incubated with puromycin (1 µM; 30 min), and global protein synthesis was measured by Western blotting using anti-puromycin antibody. ****p < 0.0001. ANOVA

To evaluate the impact of MDK signaling on protein synthesis, we used the cell-permeable, alkyne-containing puromycin analog O-propargyl-puromycin (OPP) to measure nascent protein production. After MDK antibody treatment, MB cells were incubated with OPP for 90 min and processed for direct fluorescent labeling via click chemistry using 5-FAM azide. Confocal microscopy showed that MDK suppression significantly reduced 5-FAM fluorescence, indicating decreased nascent protein synthesis (**Fig. 5I-L**). We further examined global translation by treating MDK-silenced or antibody-treated cells with puromycin followed by Western blotting with anti-puromycin antibody (SUnSET assay). As expected, MDK suppression reduced global protein synthesis in a dose-dependent manner in both Daoy and D556 MB cells (**Fig. 5M,N**). Cycloheximide served as a positive control and effectively blocked new protein synthesis. Collectively, these findings demonstrate that MDK suppression inhibits ERK and mTOR signaling cascades, leading to reduced ribosome biogenesis, decreased rRNA synthesis, and attenuated new protein production in MB cells.

### MDK suppression activates the cGAS-STING/IFN signaling pathway

Transcriptomic analyses across siRNA-treated, antibody-treated, and multiple MB cell lines consistently showed that MDK suppression activates inflammatory pathways, particularly IFN-α/β signaling (**Fig. 6A**). Heatmap visualization revealed increased expression of IFNα genes in both MDK-silenced and antibody treated conditions (**Fig. 6B**). Type I IFN signaling exerts both cell-intrinsic and cell-extrinsic antitumor effects[41]. Cell-intrinsically, IFN-α activates the IFNAR–JAK1–TYK2–STAT1/STAT2 pathway within the same cell to induce hundreds of interferon-stimulated genes (ISGs), which suppress cell proliferation, promote apoptosis, enhance MHC-I antigen presentation, and establish an antiviral state. Our RT-qPCR results confirmed that several cell-intrinsic ISGs, including OAS1, IFIT2, and ISG15, as well as cell-extrinsic genes, were significantly upregulated following MDK suppression (**Fig. 6C**). Further, upregulation of IFNα genes was further validated in additional MB cells, such as D556 and HD-MB03 (**Fig. 6D-F**). The cGAS–STING pathway is a central inducer of type I interferons (IFN-α/β) and inflammatory cytokines driving cell death through both cell-intrinsic and cell-extrinsic mechanisms[42]. ERK and AKT signaling can suppress cGAS expression and activation. Based on these observations, we hypothesized that MDK promotes MB survival by attenuating IFN signaling, suppressing pro-apoptotic ISG activation, thereby contributing to immune evasion.

**Fig. 6.**
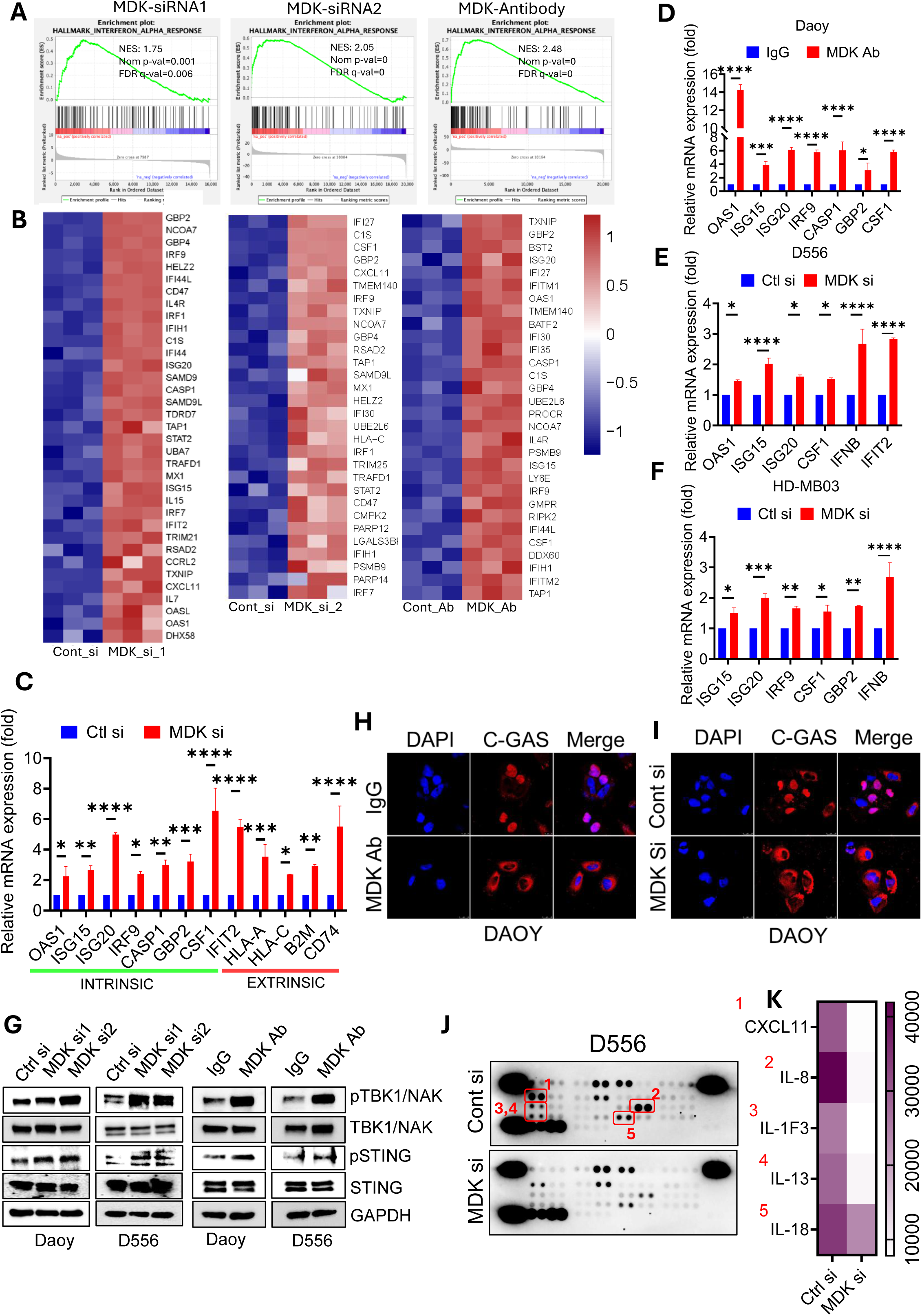
MDK suppression activates the cGAS–STING–IFN pathway in MB cells. A, GSEA plots derived from RNA-seq data showing positive enrichment of IFN-α signaling in Daoy cells transfected with two different siRNAs or treated with MDK-Ab. B, Heatmap showing upregulation of IFN-α gene expression in MDK-siRNA- or MDK-Ab-treated cells. C-F, RT-qPCR showing upregulation of cGAS/STING-IFN pathway genes in Daoy MDK-siRNA-transfected cells (C), MDK-Ab-treated cells (D), D556 MDK-siRNA cells (E), and HD-MB03 MDK-siRNA cells (F). G, Western blot showing increased p-STING and p-TBK1 in Daoy and D556 MDK-siRNA-treated or MDK-Ab-treated cells. H-I, Daoy cells that were treated with vehicle or MDK-Ab (2.5 µg; 48 h) or transfected with control or MDK siRNA, were fixed, stained with cGAS antibody, and imaged by confocal microscopy. J-K, MDK was silenced using siRNA, and lysates from control and MDK-KD cells were subjected to R&D Human Cytokine Arrays (J), with quantification shown in (K). *p < 0.05; **p < 0.01; ***p < 0.001; ****p < 0.0001.

To determine MDK regulation of cGAS-STING signaling, we examined cGAS–STING pathway activation by Western blotting. Our results showed that both MDK silencing with two independent siRNAs and MDK-antibody treatment increased cGAS–STING pathway activation in Daoy and D556 MB cells, such as increased STING, and TBK1 expression (**Fig. 6G**). Under basal conditions, cGAS is predominantly nuclear and tightly tethered to chromatin, maintaining it in an autoinhibited state. In response to cellular stress or DNA damage, cGAS can escape from chromatin and redistribute into the cytosol, where it activates STING, and produce type 1 IFNs. Thus, cytosolic relocalization is a critical licensing step that converts cGAS from a quiescent nuclear sensor into an active initiator of innate immune signaling. Since AKT signaling is known to suppress this cytosolic relocalization[43], we examined the localization of cGAS following MDK silencing or antibody treatment. Our immunofluorescence studies showed that cGAS is redistributed to the cytosol in both MDK-silenced and MDK-antibody-treated cells (**Fig. 6H-I**). Since the GAS–STING/IFN pathway induces cell death through both cell-intrinsic and cell-extrinsic mechanisms, we next examined the impact of MDK signaling on cytokine production using human cytokine arrays. Interestingly, cytokine arrays further demonstrated that MDK silencing decreased the production of pro-tumorigenic inflammatory and immunosuppressive cytokines such as IL8, IL13, IL18, and CXCL11 in MB cells (**Fig. 6J-K**). Taken together, these findings suggest that MDK suppression reactivates the cGAS–STING–IFN signaling axis, thereby promoting IFN-mediated cytotoxicity and inducing cell death in MB cells.

Given that MDK expression is heavily enriched within the malignant compartment, we utilized published single-cell RNA-sequencing datasets for CellChat analysis to infer the directionality and strength of MDK pathway interactions within the tumor microenvironment. Consistent with MDK-driven extrinsic signaling in the MB tumor microenvironment, intercellular communication modeling revealed a robust MDK signaling network in which tumor cells serve as the primary source of MDK ligands, driving dense signaling interactions toward multiple stromal cell populations (**Supplementary Fig. 1B**). Collectively, these data indicate that MB tumor cells are the primary producers of MDK and utilize MDK signaling to engage in extensive crosstalk with the surrounding microenvironment.

### MDK knockout suppresses orthotopic MB xenograft growth and enhances mice survival

To study the essential role of MDK in MB progression *in vivo*, we generated MDK-knockout (MDK-KO) cells using CRISPR gRNA transduction (**Fig. 7A**) and implanted them via intracerebellar injections. Tumor growth was monitored using the Xenogen IVIS imaging system, and survival was recorded when mice exhibited neurological symptoms. MDK-KO cells formed significantly smaller intracerebellar tumors compared with control gRNA-expressing cells (**Fig. 7B-C**) and Kaplan–Meier analysis revealed a marked extension of survival in mice bearing MDK-KO xenografts (**Fig. 7D**). Next, we examined the expression of the proliferation marker Ki67 in tumors from control and MDK-KO groups using immunohistochemistry. The IHC results showed that the number of Ki67-positive proliferating cells was significantly lower in MDK-KO tumors compared to control tumors (**Fig. 7E**). In addition, we assessed the expression of phosphorylated ribosomal protein pS6 in tumor sections. As shown in **Fig. 7F**, pS6 expression was significantly downregulated in MDK-KO Daoy tumors compared with control gRNA-expressing tumors. Collectively, these in vivo data provide genetic validation that MDK is a critical driver of MB tumor growth.

**Fig. 7.**
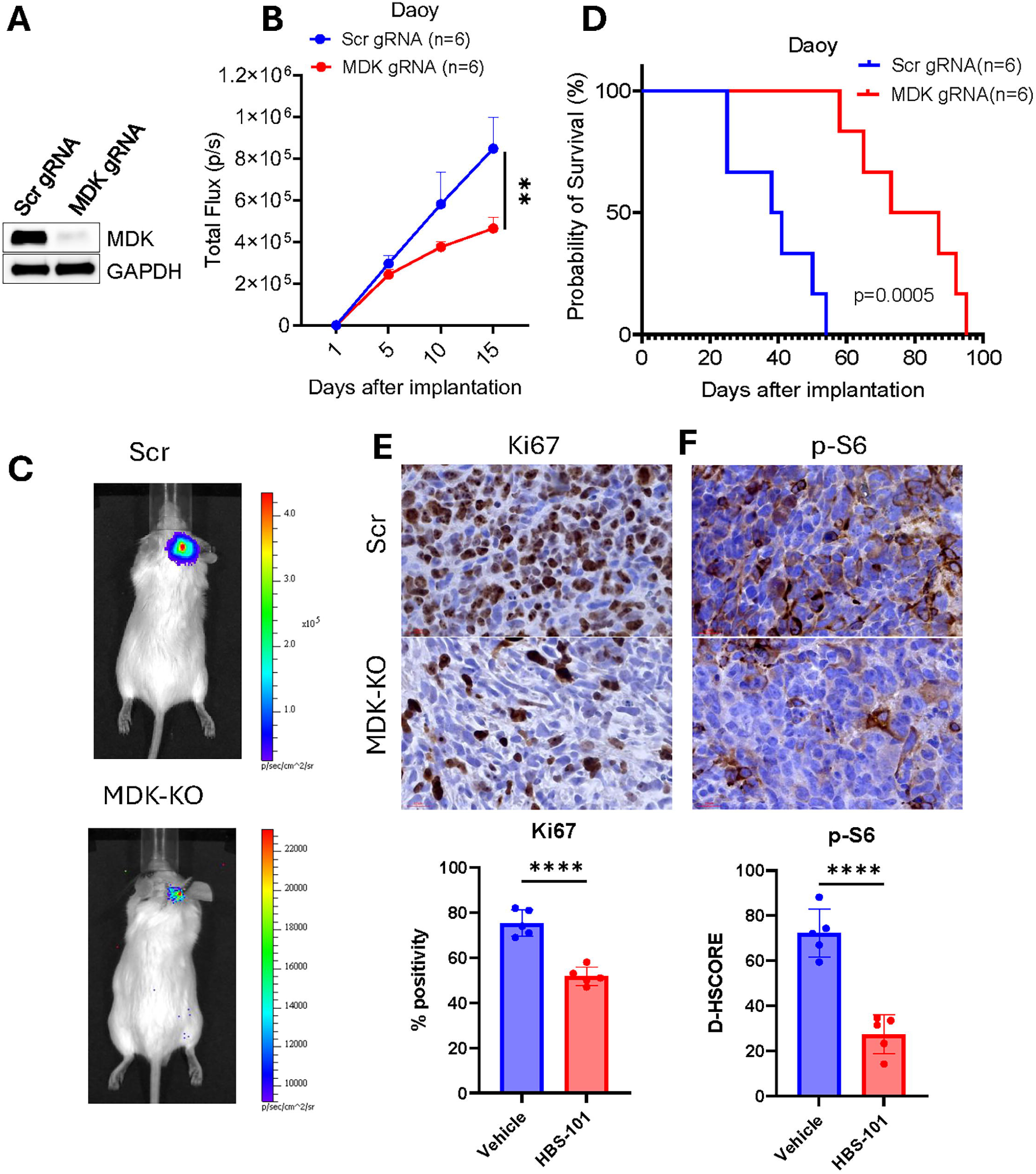
MDK is essential for MB tumor growth in vivo. A, Daoy-luc cells with MDK knockout generated using CRISPR/Cas9; loss of MDK expression was confirmed by Western blotting. B, Daoy-Scr and Daoy-MDK-KO cells (1×10⁶) were implanted into the cerebellum of female SCID mice, and tumor growth was monitored using Xenogen IVIS. C, Representative images of tumor-bearing mice at day 15 after tumor establishment. D, Kaplan–Meier survival curve comparing Daoy-Scr and Daoy-MDK-KO tumor-bearing mice. E–F, Xenograft tumor sections were analyzed for Ki67 (E) and p-S6 (F) expression by IHC. **p < 0.01, ****p<0.0001 (ANOVA).

## Discussion

Midkine (MDK) is a multifunctional heparin-binding growth factor implicated in diverse physiological and pathological processes, including several hallmark cancer traits[13, 20, 44–50]. MDK is one of the most highly expressed genes in medulloblastoma (MB)[22] and its elevated expression correlates with poor prognosis [51]. Despite this strong correlative evidence, the functional contribution of MDK signaling to MB biology has remained largely unresolved. In this study, we provide the first mechanistic and therapeutic evidence that MDK is a critical regulator of MB pathogenesis. We show that MDK signals through multiple receptors, activates key oncogenic pathways including MAPK and AKT, promotes dysregulated ribosome biogenesis and protein translation, and suppresses intrinsic IFN signaling and proinflammatory cytokine responses. Importantly, MDK knockout markedly reduced tumor growth and improved survival in orthotopic MB models. Together, these findings establish MDK as both a mechanistic driver and a therapeutic vulnerability in MB.

Independent evidence from recent single-cell, spatial transcriptomics, and cell-cell communication studies strongly supports a central role for MDK in MB. MDK is highly expressed across SHH, Group 3, and Group 4 MBs, and SAGE studies identified MDK as one of the most abundant transcripts in MB tissues, with little to no expression in normal cerebellum[22]. Single-cell analyses further reveal that MDK expression is associated with poor outcomes in SHH-α MBs enriched for Li-Fraumeni syndrome (LFS) and that MDK drives tumor-intrinsic communication networks[51]. Across SHH, Group 3, and Group 4 MBs, MDK signaling consistently emerges as a dominant communication pathway facilitating interactions between tumor cell populations[30]. In high-risk MBs, differentiating and cycling tumor cells predominantly secrete MDK, which binds to NCL on non-cycling progenitor-like cells, indicating a conserved ligand–receptor mechanism across aggressive MB subtypes. Collectively, these studies position MDK as a central regulator of intratumoral signaling in MB and reinforce its potential as a therapeutically actionable vulnerability. The results from this study corroborate these observations and provide the functional evidence that MDK promotes MB growth through activation of MAPK- and AKT-dependent ribosomal biogenesis and suppression of cGAS–STING–IFN cytotoxic signaling.

MDK exerts its biological effects through binding to a heterogeneous network of receptors that regulate survival, proliferation, migration, immune modulation, and metabolic reprogramming[13]. Unlike classical growth factors that signal through a single high-affinity receptor, MDK engages multiple receptor complexes often requiring heparan-sulfate proteoglycans such as syndecans to potentiate receptor interactions and signal propagation[29]. Multi-omic studies have shown that MDK cognate receptors including NCL, SDC2, PTPRZ1, and LRP1 are expressed in MB tumor cells and mediate intratumoral communication[30, 51, 52]. Consistent with these findings, our results confirmed expression of these receptors in MB cells. Because MDK activates multiple intracellular pathways through several receptors, targeting MDK directly may provide broader therapeutic impact than inhibiting each single receptor or pathway.

A key mechanistic insight emerging from our study is that MDK-mediated activation of ERK and AKT/mTOR establishes a signaling architecture that converges on MYC, a central oncogenic driver in MB. ERK and mTOR stabilize and activate MYC through phosphorylation-dependent post-translational modifications, enhancing MYC transcriptional regulation of ribosomal proteins, rRNA transcription, aminoacyl-tRNA synthetases, and translation-initiation factors[53],[54]. MYC-driven ribosome biogenesis is further reinforced by a MYC–mTOR feed-forward loop, where MYC transcriptionally upregulates mTOR pathway components, while mTORC1 enhances MYC protein stability and translation[37, 55–57]. ERK also supports ribosome biogenesis by activating RSK and promoting UBF-dependent rDNA transcription[58]. Our findings that MDK inhibition suppresses pERK, pAkt, pS6, p4EBP1, MYC targets, rRNA species, and global protein synthesis align with this integrated model and position MDK as a master upstream regulator of the MYC–mTOR translational axis in MB.

MDK functions as a soluble growth factor that acts through both autocrine and paracrine mechanisms within the tumor microenvironment (TME)[29]. These dual modes allow MDK to influence not only tumor-intrinsic pathways but also stromal, immune, and vascular compartments that support tumor progression. MDK is emerging as a potent regulator of TME immunosuppression: it promotes Treg expansion, suppresses CD8⁺ T-cell cytotoxicity, drives M2-like macrophage polarization, impairs NK-cell function via soluble MICA/B, and reprograms dendritic cells toward tolerogenic states through STAT3 activation[10, 29, 59–62]. High MDK expression correlates with immune exclusion, poor outcome, and reduced response to immunotherapy, while MDK inhibition enhances antitumor immunity and improves responses to CAR-based and checkpoint therapies[63, 64]. Our RNA-seq analyses across MDK-silenced and antibody-treated MB cells showed increased activation of immune and inflammatory pathways, including strong induction of IFN-α signaling and engagement of the cGAS–STING pathway. The cGAS–STING axis is a central cytosolic DNA-sensing mechanism that drives type I IFN (IFN-α/β) and inflammatory cytokine production to induce cell death via both cell-intrinsic and extrinsic mechanisms[42]. ERK and AKT signaling are known to suppress cGAS transcription and activation, and AKT can inhibit cytosolic relocalization of cGAS by stabilizing chromatin tethering[43, 65, 66]. Consistent with this biology, our results using Western blotting and immunofluorescence showed that MDK suppression increased cGAS, STING, and TBK1 activation and promoted cytosolic redistribution of cGAS an essential licensing step for STING activation and type I IFN production. Type I IFN signaling exerts both cell-intrinsic and extrinsic antitumor effects. Cell-intrinsically, IFN-α activates the IFNAR–JAK1–TYK2–STAT1/STAT2 pathway, inducing hundreds of ISGs that suppress proliferation, promote apoptosis, enhance MHC-I antigen presentation, and generate an antiviral-like state[41, 67, 68]. Our transcriptomic and functional analyses showed strong upregulation of multiple intrinsic ISGs (OASL, IFIT3, ISG15, MX1/MX2) as well as extrinsic immune genes (HLA-A/B, CXCL chemokines) following MDK inhibition, supporting enhanced antigen presentation and cytotoxic immune recruitment. Together, these findings suggest that MDK contributes to immune evasion in MB and that MDK-targeted therapy has the potential to promote both tumor-intrinsic cell death and improved antitumor immunity. Future studies using immunocompetent MB models and comprehensive TME profiling will be essential to fully define MDK’s immunomodulatory functions in vivo.

In summary, our findings identify MDK as a previously underappreciated yet central regulator of medulloblastoma biology. By integrating receptor-level signaling, oncogenic kinase activation, MYC–mTOR–ribosome control, and innate immune suppression, MDK establishes a multifaceted network that supports tumor growth, metabolic adaptation, and immune evasion. The discovery that MDK suppression produces potent antitumor effects underscores MDK as a compelling therapeutic vulnerability. Continued investigation into MDK roles in tumor heterogeneity, ribosome biology, and immune modulation will further refine its translational potential and may ultimately lead to improved outcomes for patients with high-risk MB.

## Authorship

Conceptualization: G.R.S., H.B.N., R.K.V., S.V.,; Methodology: S.J., P.P.V., J.D.J., N.A., B.S., U.P.P., S.V., P.S., M.K.R., A.J.B., ; Software and Bioinformatics: Y.C., Z.Y., S.Z., N.A., ; Writing, Review, Editing: G.R.S., R.K.V., H.B.N., K.Y., ; Supervision: G.R.S.,. All authors read and approved the manuscript.

---

**Supp Fig. 1A.**
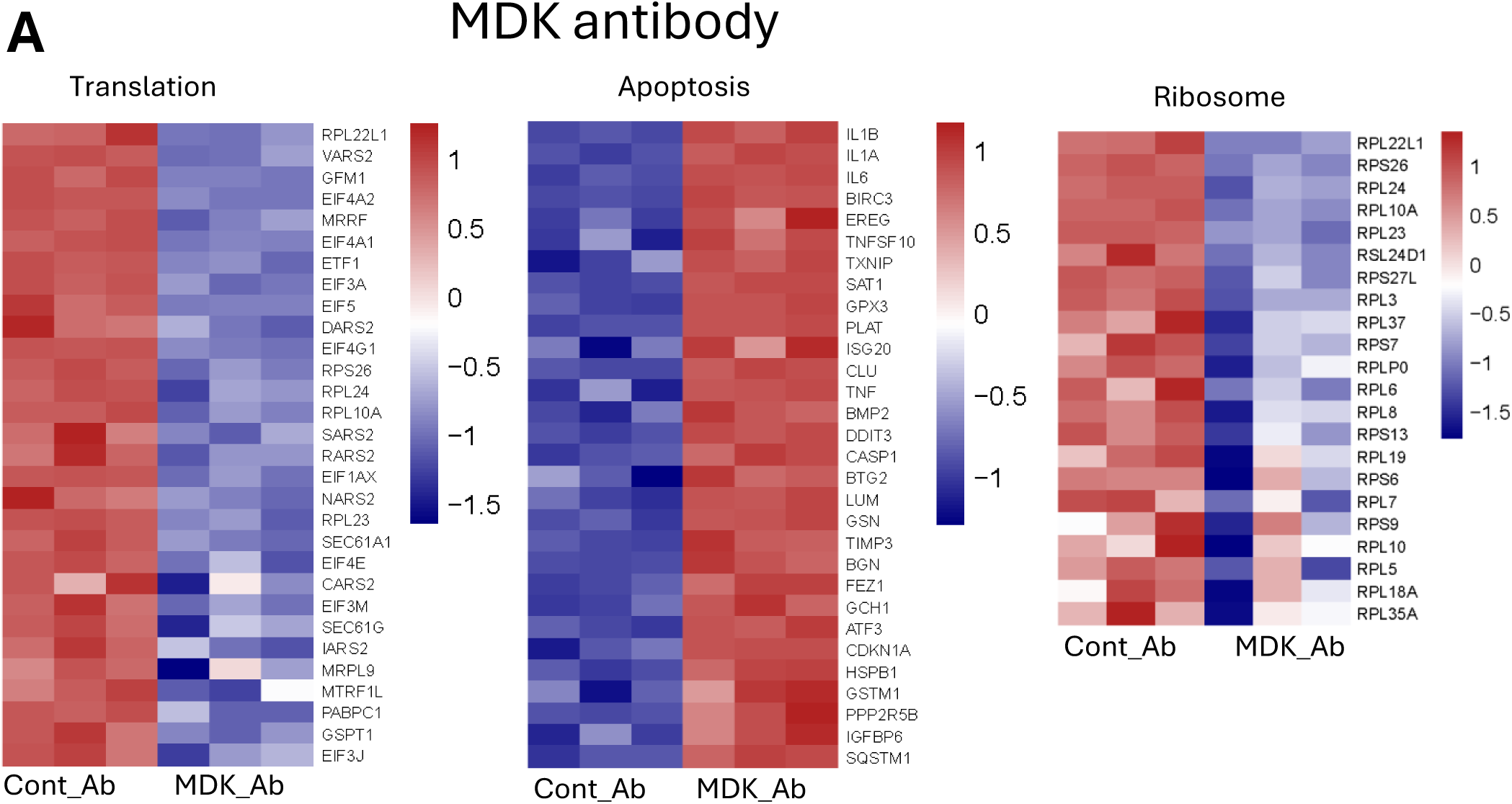
Daoy cells treated with control IgG or MDK-antagonizing antibody (48 h) were subjected to RNA-seq. Heatmap showing downregulation of ribosome/translation genes and upregulation of apoptosis-related genes.

**Supp Fig. 1B.**
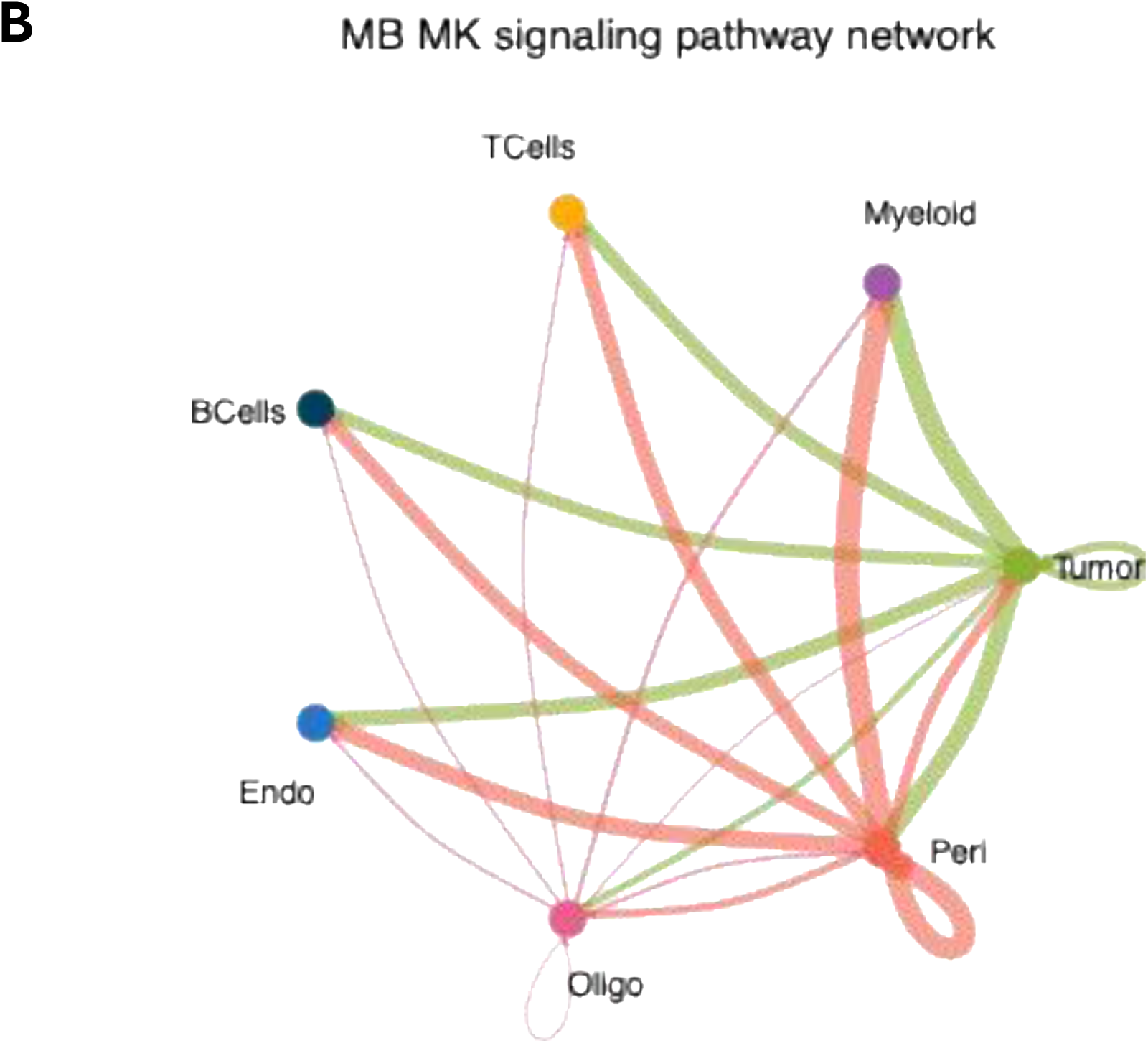
Intercellular communication network of the Midkine (MK) signaling pathway inferred via CellChat analysis.

